# Cryo-EM structures of tau filaments from SH-SY5Y cells seeded with brain extracts from cases of Alzheimer’s disease and corticobasal degeneration

**DOI:** 10.1101/2023.04.13.536725

**Authors:** Airi Tarutani, Sofia Lövestam, Xianjun Zhang, Abhay Kotecha, Andrew C. Robinson, David M.A. Mann, Yuko Saito, Shigeo Murayama, Taisuke Tomita, Michel Goedert, Sjors H.W. Scheres, Masato Hasegawa

**Author notes:** These authors contributed equally (Airi Tarutani, Sofia Lövestam).

## Abstract

The formation of amyloid filaments through templated seeding is believed to underlie the propagation of pathology in most human neurodegenerative diseases. A widely used model system to study this process is to seed amyloid filament formation in cultured cells using human brain extracts. Here, we report the electron cryo-microscopy (cryo-EM) structures of tau filaments from undifferentiated seeded SH-SY5Y cells, that transiently expressed N-terminally HA-tagged 1N3R or 1N4R human tau, using brain extracts from individuals with AD or CBD. Although the resulting filament structures differed from those of the brain seeds, some degree of structural templating was observed. Studying templated seeding in cultured cells, and determining the structures of the resulting filaments, can thus provide insights into the cellular aspects underlying neurodegenerative diseases.

## Introduction

The formation of abundant filamentous tau inclusions is a defining characteristic of a number of neurodegenerative diseases, collectively referred to as tauopathies (1). The identification of disease-causing mutations in *MAPT*, the tau gene, in dominantly inherited forms of frontotemporal dementia, together with the invariable presence of abundant filamentous tau inclusions, established that the formation of tau filaments is sufficient to cause neurodegeneration and dementia.

Six tau isoforms are expressed by alternative mRNA splicing in adult human brains (2). Based on the expression (or not) of a 31 amino acid repeat, they can be divided into three isoforms with 3 repeats each (3R) and three isoforms with 4 repeats each (4R). Disease-associated filaments are made of either 3R+4R tau, 3R tau or 4R tau. Work using electron cryo-microscopy (cryo-EM) identified two distinct folds for 3R+4R tauopathies (3,4), one fold for 3R tauopathies (5) and five folds for 4R tauopathies (6-8). Although specific tau folds define different diseases, several conditions share a fold. Commonalities in structure suggest that similar molecular mechanisms may lead to the formation of amyloid filaments in these diseases.

Experimental evidence indicates that filamentous tau may spread through templated seeding in human brains, reminiscent of prions. Thus, assembled tau has been found to move from the outside to the inside of cultured cells (9). Moreover, in the brains of transgenic mice, tau seeds induced the local aggregation of human tau, followed by the spread of tau assemblies to distant brain regions (10). The observations that specific tau filament structures characterise diseases and that these structures are identical in different brain regions (8,11,12), support templated seeding. Two assumptions are central to the hypothesis of templated seeding. The first is that a small amount of tau filaments, the seed, amplifies filament formation beyond the levels that would have happened spontaneously. The second is that seeded tau filaments have the same structures as the seeds.

It is often assumed that seeds and seeded aggregates have identical structures. However, when their structures were determined (seeds of assembled α-synuclein from multiple system atrophy brains and seeded aggregates of recombinant α-synuclein), they were different (13). Similar experiments have not been done for tau. Here we transiently expressed N-terminally HA-tagged 1N3R and 1N4R tau in SH-SY5Y cells, which were exposed to tau seeds from the brain of an individual with either Alzheimer’s disease (AD) or corticobasal degeneration (CBD). The structures of the seeded aggregates were similar, but not identical, to those of the seeds.

## Results

### Aggregation of tau in SH-SY5Y cells seeded with tau filaments from Alzheimer’s disease or corticobasal degeneration

The cryo-EM structures of tau filaments were determined following seeded aggregation in SH-SY5Y cells using AD and CBD seeds (Figures 1-4; Supplementary Figure 1). In AD, abundant neurofibrillary lesions in occipital cortex were immunoreactive with anti-tau antibodies AT8, RD3 and Anti-4R. In CBD, abundant neuronal and glial inclusions in putamen were labelled by AT8 and RD4, but not RD3 (Supplementary Figure 2a). Immunoblotting of sarkosyl-insoluble fractions with T46 showed major tau bands of 60, 64 and 68 kDa and a minor band of 72 kDa in AD, and major bands of 64 and 68 kDa in CBD. RD3 labelled the 60, 64 and 68 kDa bands in AD, whereas RD4 visualised the 64, 68 and 72 kDa bands in AD, and the 64 and 68 kDa bands in CBD (Supplementary Figure 2b).

**Figure 1:**
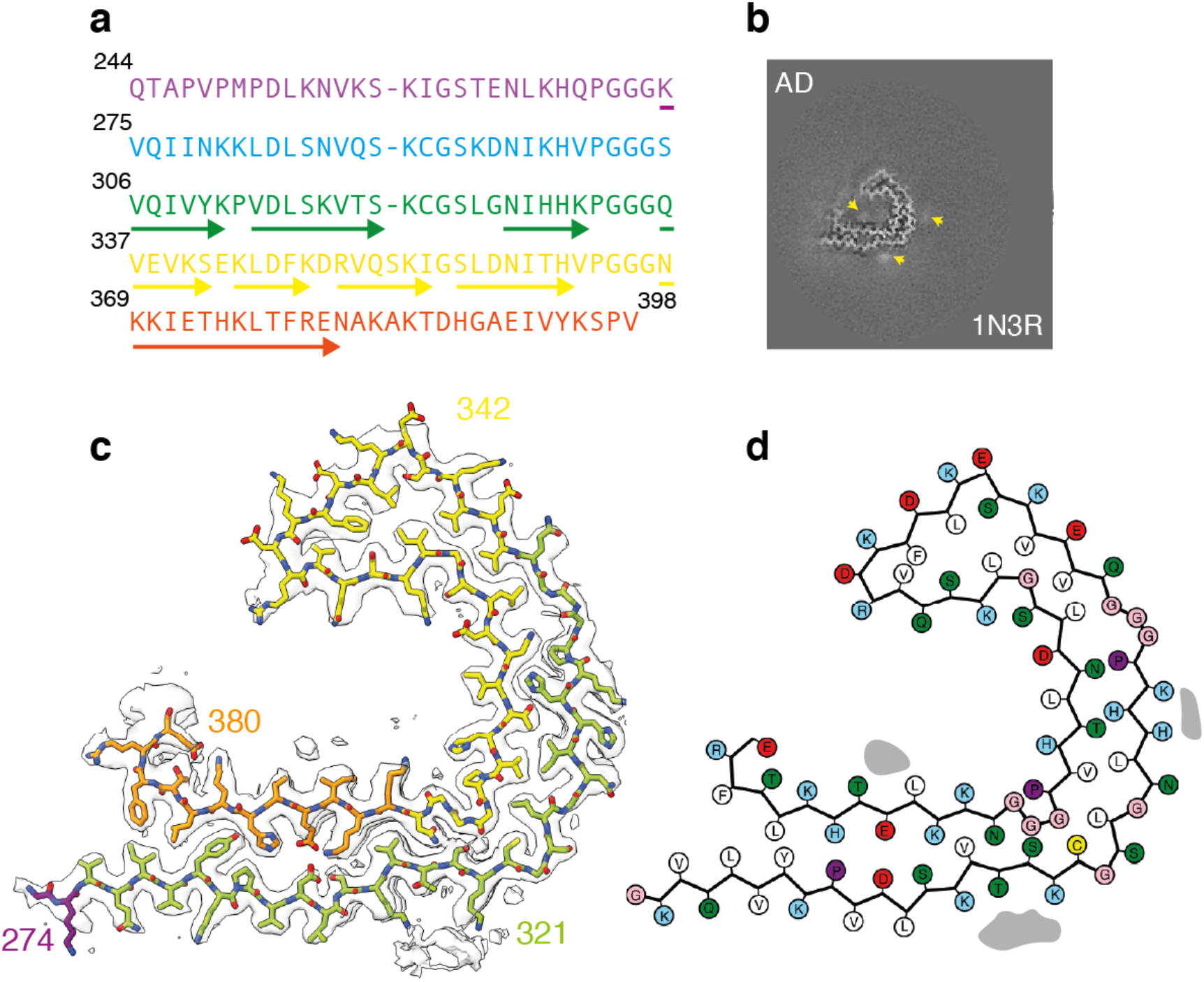
Tau structure in SH-SY5Y cells seeded with filaments from Alzheimer’s disease. a. Amino acid sequence of tau residues 244-398 coloured by repeat: R1 is purple; R2 is blue; R3 is green; R4 is yellow; the C-terminal domain is orange. b. XY-cross-section of the reconstructed density with a projected thickness of approximately 1 β-rung (4.75 Å). Extra densities are indicated with yellow arrows. c. Reconstructed density (in transparent white) and atomic model (with the same colours as in panel a). d. Schematic of the residues in the structure. Positively charged residues are shown in blue; negatively charged residues in red; polar residues in green; apolar residues in white; cysteines in yellow; glycines in pink and prolines in purple.

Sarkosyl-insoluble fractions were introduced by lipofection into SH-SY5Y cells that transiently expressed N-terminally HA-tagged 1N3R or 1N4R tau (Supplementary Figure 3a), as described (14). Immunoblot analysis of sarkosyl-insoluble fractions from seeded SH-SY5Y cells showed accumulation of HA-1N3R and HA-1N4R tau (Supplementary Figure 3b). By immunoelectron microscopy, abundant filaments made of HA-1N3R tau or HA-1N4R tau were in evidence following exposure to AD or CBD seeds (Supplementary Figure 3c).

### Structure of HA-1N3R tau filaments seeded by extract of occipital cortex from Alzheimer’s disease

Following seeding with extract from the occipital cortex of a patient with AD, we observed a single filament type in sarkosyl-insoluble fractions of SH-SY5Y cells that transiently expressed HA-1N3R tau. We calculated a cryo-EM 3D reconstruction of this filament to 2.5 Å resolution. The structure revealed the presence of a single protofilament with a C-shaped fold (Figure 1). The ordered core comprises residues K274-E380 of 3R tau and consists of one residue of R1, the whole of R3, the whole of R4 and 12 amino acids of the C-terminal domain of tau. The protofilament, which comprises 8 β-strands, is similar to the Alzheimer tau fold of paired helical filaments (PHFs) and straight filaments (SFs) (3,11). The r.m.s.d. between the seeded structure and PDB entry 6hre is 2.41 Å. Besides the fact that PHFs and SFs are always made of two identical protofilaments, the single protofilament of the seeded structure has two minor conformational differences with the Alzheimer tau fold (Figure 2).

**Figure 2:**
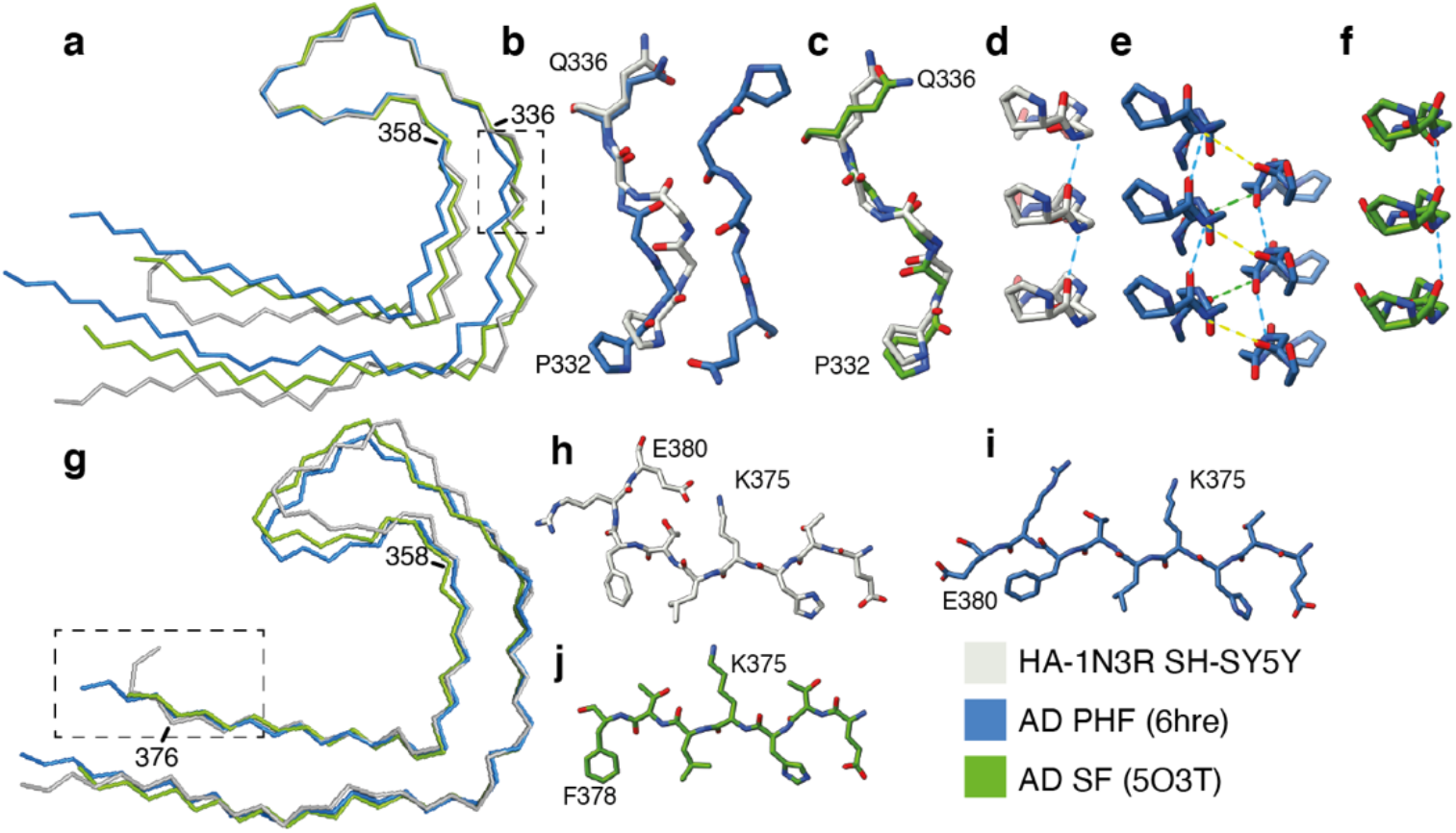
Comparison of the seeded structure and the structure of the seeds from Alzheimer’s disease. a. Overlay of main-chain traces of the seeded structure (grey) and the PHF (blue) and SF (green) from Alzheimer’s disease (AD), aligned on the tip of the C-shaped fold (residues 336-358). b-f. Detailed views of the ^332^PGGGQ^336^ motif with the same colours as in panel a. Intra-protofilament hydrogen bonds are shown with blue dashed lines and inter-protofilament hydrogen bonds with yellow and green dashed lines. Panels b and c are top views of the comparison between the seeded structure and the PHF and SF, respectively. Panels d-f are side views of the seeded structure, the PHF and the SF, respectively. g. Overlay of main chain traces of the seeded structure, the PHF and the SF, aligned on the ends of the C-shaped fold (residues 358-376). h-j. Detailed views of the C-terminal part of the ordered cores of the seeded structure, the PHF and the SF, respectively.

The first difference is in the conformation of the ^332^PGGG^335^ motif. In PHFs, there are inter-protofilament hydrogen bonds between the backbones of G333 and G334, and between the side chain of Q336 and the backbone of K331. In the structure of seeded aggregates, in the absence of a second protofilament, residues ^332^PGGG^335^ adopt a different conformation, with the main chain carbonyl of P332 and the main chain nitrogen atom of G333 forming hydrogen bonds within the protofilament along the helical axis instead. This alternative conformation leads to an opening of the C-shaped fold compared to the AD fold. Moreover, there is an additional density in front of H329 and K331 in the seeded structure that is not found in PHFs. The molecules that form this density are probably negatively charged, so that they can counter the positive charges of the side chains of H329 and K331. In PHFs, K331 orients towards the side chain of E338 of the opposing protofilament. Additional densities, in front of K317, K321 and β8 in the inside of the C-shaped protofilament, are found in the tau folds of both seeded aggregates and AD. In this region, the seeded structure is more similar to AD SFs than to PHFs. AD SFs also have intra-protofilament hydrogen bonds at P331 and G322, an additional density in front of H329 and K331 and a more open C-shaped fold. The seeded structure has an overall r.m.s.d. of 1.79 Å with AD SFs.

The second difference is in the C-terminal part of the ordered core. Residues R379 and E380, which form part of β8 in the Alzheimer fold (11), turn away from the opposing β1 in the seeded structure, with the side chain of E380 forming a salt bridge with the side chain of K376.

### Structure of HA-1N4R tau filaments seeded by extract from putamen of corticobasal degeneration

Following seeding with extract from the putamen of a patient with CBD, we initially observed a single filament type in sarkosyl-insoluble fractions of SH-SY5Y cells that transiently expressed HA-1N4R tau. An initial 3D reconstruction revealed a four-layered fold with 9 β-strands (β1-β9). Smeared density around the C-terminus of the ordered core indicated that the reconstruction was calculated from a mixture of different structures. Two filament types were resolved by 3D classification, for which we obtained resolutions of 2.5 and 2.3 Å (Figure 3). They are identical between residues 277 and 360, but filament type 1 has an additional 15 ordered residues at the C-terminus. It extends from 363-377. Residues 364-367 form a hairpin, whereas residues 372-375 form a tenth β-strand (β10) that packs against β9 (Figure 3). In type 2 filaments, these residues are disordered.

**Figure 3:**
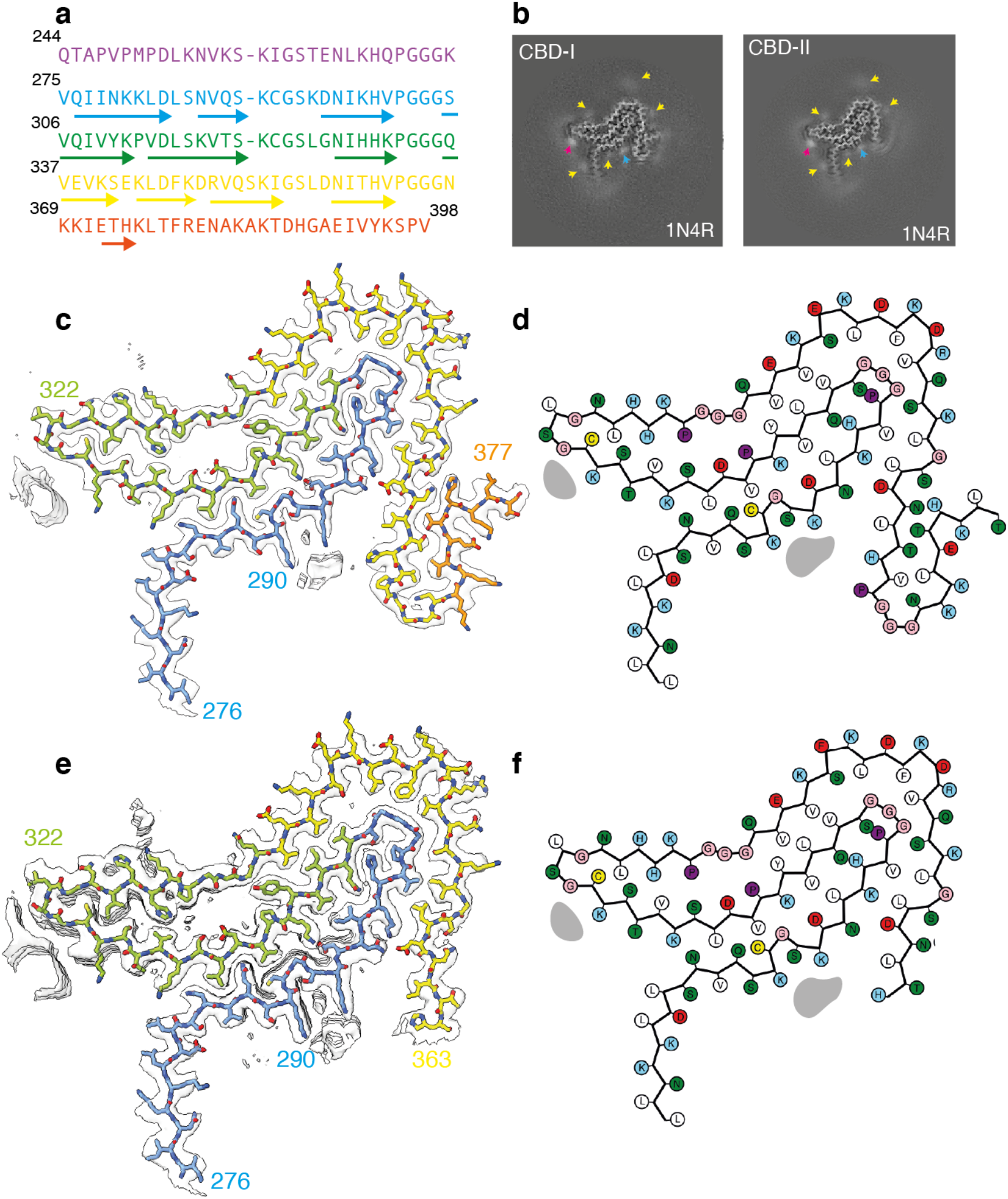
Tau structure in SH-SY5Y cells seeded with filaments from corticobasal degeneration. a. Amino acid sequence of tau residues 244-398. b. XY-cross-section of the reconstructed density with a projected thickness of approximately 1 β-rung (4.75 Å). Extra densities are indicated with yellow, pink and blue arrows. c. Reconstructed density and atomic model. d. Schematic of the residues in the structure. Colours are as in Figure 1.

The core shared between both filament types adopts a four-layered fold that is similar to that of CBD (6,7). Residues 277-362 have almost identical conformations, with an r.m.s.d. of 1.96 Å between the seeded structures and CBD filaments (Figure 4). The main differences are located in the amino- and carboxy-terminal regions. In the carboxy-terminal part of the CBD fold, residues 364-367 are in an extended conformation, whereas amino acids 368-380 form a long β-strand (β11). The latter wraps around an internal density that is coordinated by the side chains of K290 and K294 on one side of a large internal cavity and K370 on the other side. In the seeded structures, residues 364-367 form a hairpin or are disordered. As a result, there is no internal cavity and an additional density is only coordinated by the side chains of K290 and K294. In the amino-terminal part of the CBD fold, residues 274-280 form a β-strand that packs against the carboxy-terminal half of β11, closing the cavity. D283 forms part of a sharp turn in the CBD fold and its side chain points outwards. In the seeded structures, D283 is part of an extended β-strand that does not pack with another strand and the side chain of D283 points towards the protofilament core. The additional densities in both folds are also different in size. In the plane perpendicular to the helical axis, the additional density in the CBD fold has an elongated shape and measures 9 x 4 Å. In the seeded structures, the additional density is smaller and less elongated, measuring approximately 5 x 4 Å.

**Figure 4:**
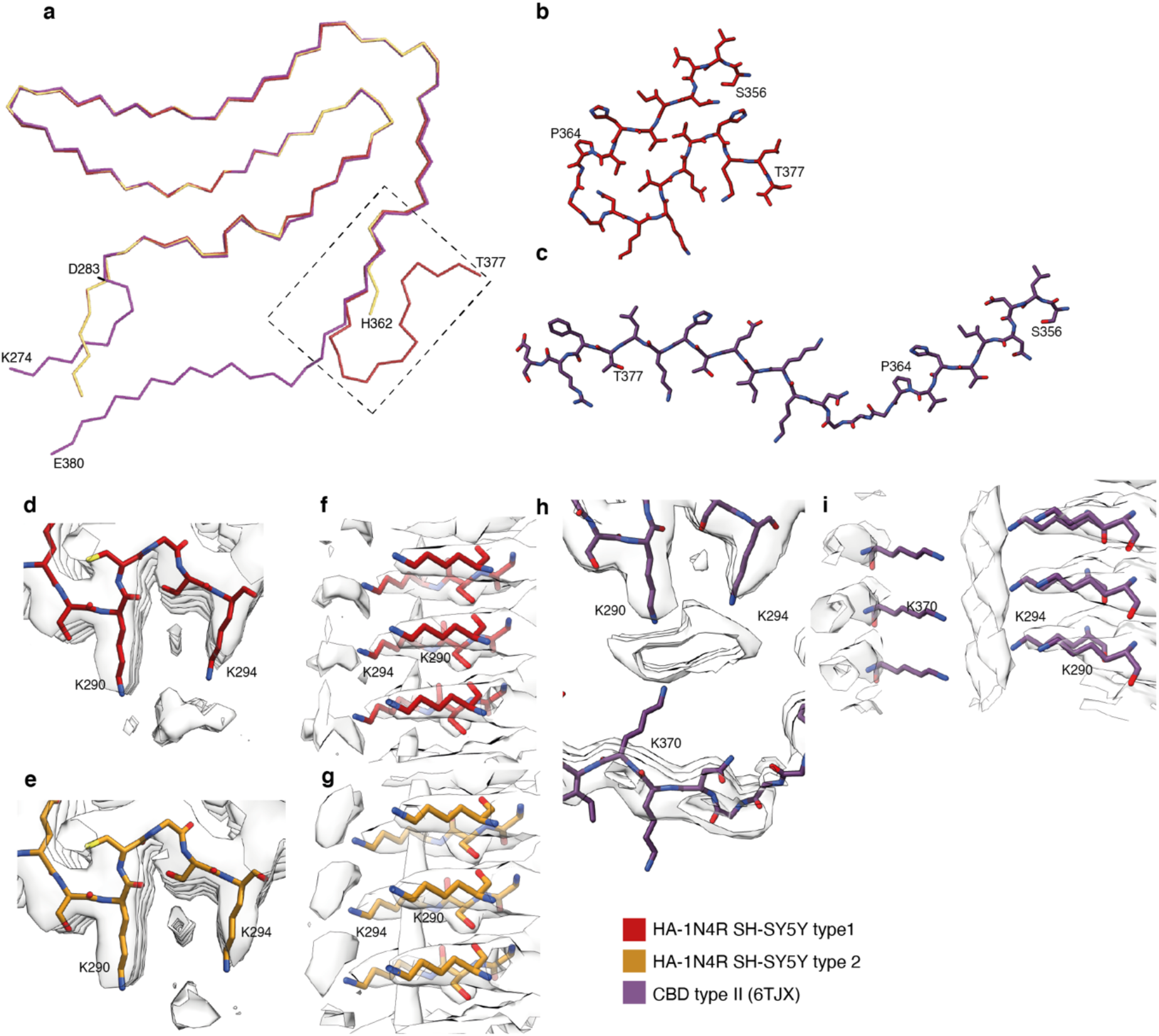
Comparison of the seeded structure and the structure of the seeds from corticobasal degeneration. a. Overlay of main-chain traces of the seeded structures (type 1 in red; type 2 in orange) and the type II filaments from corticobasal degeneration (CBD in purple) aligned on residues 283-362. b. Detailed view of residues 356-377 in the type 1 seeded structure. c. Detailed view of residues 356-380 in the CBD Type II filament. d-e. Top and side views of the extra density in the type 1 seeded structure. f-g. Top and side views of the extra density in the type 2 seeded structure. h-i. Top and side views of the extra density in the CBD type II filament.

## Discussion

Following seeding with the sarkosyl-insoluble fraction from AD brain, abundant filaments of HA-1N3R tau with a single protofilament were present that closely resembled the Alzheimer tau fold. As reported previously (14), more seeded tau aggregates formed following expression of HA-1N3R than HA-1N4R tau. Tau filaments from AD, prion protein amyloidoses, familial British dementia, familial Danish dementia and primary age-related tauopathy are made almost entirely of R3, R4 and 10-12 amino acids after R4, all of which are sequences that 3R and 4R tau have in common (3,8,11,12,15). It is therefore not surprising that tau filaments from the brains of individuals with AD are made of all six tau isoforms (16). It remains unclear why seeding of SH-SY5Y cells with the sarkosyl-insoluble fraction from AD brain is more efficient with expression of HA-1N3R tau than HA-1N4R tau.

Our findings with seeds from AD and CBD brains raise the question of why seeded tau aggregates were made of a single rather than two identical tau protofilaments. In extracts from AD brains, only filaments with two protofilaments have been observed. In extracts from human CBD brains, both filaments with a single and a double protofilament have been observed. Previous work using recombinant amino and carboxy-truncated tau fragments has shown that they assembled into AD PHFs with two identical protofilaments. Single protofilaments with the AD fold were observed following the extension and pseudo-phosphorylation of the fragments at the carboxy-terminus (17). It remains to be determined if the carboxy-terminal region of tau, or the presence of only the 1N3R isoform, is inhibitory towards the formation of protofilament doublets in human brains. It also remains to be shown if PHFs, SFs or CBD filaments with two protofilaments made of full-length tau form by the gradual merging of two protofilaments over time.

Following seeding with the sarkosyl-insoluble fraction from CBD brain, we observed two types of filaments that were made of a single protofilament of HA-1N4R tau that resembled the CBD tau fold. The two filament types differed in the carboxy-terminal regions of their ordered cores. Seeded aggregates were not observed upon expression of HA-1N3R tau, which is in agreement with the observation that 3R tau monomers cannot be incorporated into the CBD fold (7). The main differences between the CBD fold and those of the seeded aggregates were in the region flanking a large internal cavity in the CBD fold that contains an additional density of unknown molecular composition. In seeded aggregates, there was no cavity and the additional density was smaller than in CBD. It therefore appears likely that the additional densities in the seeded structures and the CBD fold are made of different molecules. It remains to be established if the shape of the amyloid fold and its cavity determine the size of the density or if it is the density that determines the shape of the amyloid fold and its cavity.

In conclusion, we show that seeded aggregation of SH-SY5Y cells that express HA-tagged 1N3R or 1N4R tau, with extracts of AD or CBD brains, yields structures that resemble, but are not identical to, the seed structures. Thus, similar to our previous observations for the *in vitro* assembly of purified, recombinant α-synuclein (13), one cannot assume that the seeded amyloid structures are correctly replicated in cell-based seeded aggregation experiments. Whether it is relevant that the correct structures are replicated will depend on the scope of the seeded aggregation experiments. Still, our observations suggest that some degree of templating is taking place when SH-SY5Y cells are seeded with brain extracts of AD or CBD brains. Thereby, these types of experiments hold the potential to inform on cellular aspects of disease. Since specific filament structures define different diseases, investigating which cell types replicate the same structures observed in disease could help to identify the cell types that are important in disease. AD is characterised by abundant tau inclusions in nerve cells, the formation of which may begin in the entorhinal cortex (18). CBD is characterised by abundant neuronal and glial tau inclusions, with astrocytic plaques being pathognomonic (19). It has been suggested that CBD may begin as an astrogliopathy, with neuronal inclusions becoming the predominant lesion type in advanced disease (20). The undifferentiated SH-SY5Y cells used in this study may be imperfect models of any type of brain cells. Therefore, future experiments should investigate the effects of seeding with brain extracts from AD, CBD and other diseases on the structures of seeded filaments on a variety of cell types, including neurons, glial cells, and differentiated SH-SY5Y cells.

## Materials and Methods

### Tau seeds

AD was in a 65-year-old female from the U.K. who died with a clinicopathological diagnosis following a 9-year history of progressive dementia. This case has not been described previously. CBD was in a 74-year-old female from Japan that has been described before [case 1 in (7)]. Sarkosyl-insoluble fractions were prepared from occipital cortex of the AD case and putamen of the CBD case, as described (21). Briefly, 0.25 g of brain tissue was homogenised in 20 vol (w/v) extraction buffer (10 mM Tris-HCl, pH 7.5, containing 10% sucrose, 0.8 M NaCl, 1 mM EGTA). Homogenates were brought to 2% sarkosyl and incubated for 30 min at 37° C. Following a 10 min centrifugation at 27,000g, the supernatants were spun at 113,000g for 20 min at room temperature. The pellets were resuspended in 500 μl extraction buffer and spun again at 113,000g for 20 min. Sarkosyl-insoluble pellets were the tau seeds; they were resuspended in 640 μl/g saline, sonicated 10 times for 1 s each at power output 4 (TAITEC VP-5s) and used for lipofection into SH-SY5Y cells and immunoblotting.

### Cell culture

Human neuroblastoma SH-SY5Y cells (22) were cultured at 37° C and 95%O_2_/5%CO_2_ in Dulbecco’s modified Eagle/F-12 medium (DMEM/F-12, Sigma Aldrich), supplemented with 10% foetal calf serum, 1% penicillin-streptomycin glutamine (Gibco) and 1% MEM non-essential amino acid solution (Gibco). They were plated at 3 x 10^5^ cells/well and grown to 60-70% confluence in collagen-coated six-well plates, prior to transfection with 2 μg pcDNA3.1 encoding N-terminally HA-tagged human 1N3R tau or 1N4R tau (14) using X-tremeGENE 9 (Roche Life Sciences), following the manufacturer’s instructions. Cells were incubated for 8 h at 37° C, before introduction of 2 μl seeds (diluted 1:5) using MultiFectam (Promega), following the manufacturer’s instructions. Transiently transfected and seeded SH-SY5Y cells were grown for 3 days at 37° C.

### Seeded tau filaments from SH-SY5Y cells

SH-SY5Y cells were washed with PBS and harvested with 1 ml extraction buffer containing 1% sarkosyl. Cell extracts were then sonicated on ice for 15 s using 5 cycles of 3 s on and 3 s off (Seiko Biomic 7040 Ultrasonic Processor). After a 30 min incubation at 37° C, the cell extracts were spun at 113,000g at room temperature for 20 min. Supernatants served as the sarkosyl-soluble fractions; pellets were washed in 30 mM Tris-HCl, pH 7.5, and spun again at 113,000g for 20 min. These sarkosyl-insoluble pellets were the seeded tau aggregates; they were resuspended in 50 μl 30 mM Tris-HCl, pH 7.5, sonicated and used for immunoblotting and immunoelectron microscopy. Prior to cryo-EM, the sarkosyl-insoluble fractions were stored at -80° C.

### Antibodies

The following anti-tau antibodies were used: T46 (mouse monoclonal, specific for residues 404-441, Thermo Fisher Scientific); TauC (mouse monoclonal, specific for residues 429-441, Cosmobio); RD3 (mouse monoclonal, specific for 209-224, Millipore); RD4 (mouse monoclonal, specific for 275-291, Millipore); anti-4R (rabbit polyclonal, specific for 275-291); AT8 (mouse monoclonal, specific for pS202 and pT205, Thermo Fisher Scientific); pS396 (mouse monoclonal, specific for pS396, Calbiochem). Monoclonal and polyclonal anti-HA antibodies (HA3663 and H6908) were purchased from Sigma Fine Chemicals.

### Immunoblotting and immunohistochemistry

Immunoblotting was performed as described (21). Sarkosyl-insoluble and sarkosyl-soluble fractions were added to SDS-sample buffer and boiled for 3 min. They were separated on 4-20% polyacrylamide gradient gels (Wako) or on home-made 10% polyacrylamide gels (Wako). Protein concentrations were determined using a BCA kit (Thermo Fisher Scientific). For immunohistochemistry, brain tissues were fixed using 10% formalin neutral buffer solution (Wako) and embedded in paraffin. Following deparaffinisation, the sections were autoclaved in 10 mM citrate buffer (pH 6.0) at 120° C for 10 min and treated with 100% formic acid for 10 min. After washing in water, the sections were incubated with 3% hydrogen peroxide in PBS for 30 min, blocked with 10% foetal calf serum in PBS containing 0.3% Triton X-100 and incubated overnight in primary antibodies at room temperature. Following incubation with biotinylated secondary (Vector Laboratories) for 1 h, labelling was developed with diaminobenzidine for 20 min, using an avidin-biotin complex staining kit (Vector Laboratories).

### Immunoelectron microscopy

Immunoelectron microscopy was performed as described (21). Briefly, sarkosyl-insoluble fractions were dropped onto carbon-coated 300-mesh copper grids (Nissin EM). The grids were immunostained with anti-HA antibody (1:100) and anti-pS396 tau antibody (1:100). Secondary antibodies were conjugated to 5 nm gold particles (Cytodiagnostics, 1:50). Immunostained grids were negatively stained with 2% phosphotungstic acid and dried. Images were recorded using a JEOL JEM-1400 electron microscope, at a magnification of X15,000.

### Electron cryo-microscopy

Pellets from 6-well seeded plates were resuspended in 10 μl of 50 mM Tris, pH 7.4, 50 mM NaCl. Three μl were applied to glow-discharged (Edwards S150B) holey carbon grids (Quantifoil Au R1.2/1.3, 300 mesh) that were plunge-frozen in liquid ethane using Vitrobot Mark IV (Thermo Fisher Scientific) at 100% humidity and 4° C. Cryo-EM images were collected on a G4 Titan Krios electron microscope operated at 300 kV and equipped with a cold field-emission gun (CFEG), a Selectris X energy filter and a Falcon-4 direct electron detector at the Research and Development Facility of Thermo Fisher Scientific (Eindhoven, The Netherlands). Images were recorded at a dose of 40 electrons per Å^2^, using EPU software and converted to tiff format using relion_convert_to_tiff prior to processing. See Supplementary Table 1 for additional data collection parameters.

### Helical reconstruction

Datasets were processed in RELION using standard amyloid helical reconstruction procedures. Movie frames were gain corrected, aligned and dose weighted using RELION’s own motion correction program (23). Contrast transfer function (CTF) parameters were estimated using CTFFIND4-1 (24). Subsequent image-processing steps were performed using helical reconstruction methods in RELION (25). Filaments were auto-picked using the helical implementation of Topaz. Particles were extracted in box sizes of 1028 or 768 and down-scaled to 256 or 128 pixels. Reference-free 2D classification was used to discard suboptimal images and measure cross-over distances for initial model calculation using relion_helix_inimodel2d (26). 3D auto-refinements were performed using standard procedures in RELION, including optimisation of the helical twist and rise parameters. 3D classification without performing angular searches was used to separate the two types of HA-1N4R tau filaments obtained by seeding with extracts from putamen of CBD and to select optimal particles for final refinements. To improve the resolution of the maps, Bayesian polishing and CTF refinement were performed (27). Final maps were sharpened using post-processing procedures in RELION and their resolutions calculated based on the Fourier shell correlation (FSC) between two independently refined half-maps at 0.143 (28).

### Model building

Atomic models of tau filaments from AD- and CBD-seeded SH-SY5Y cells were built using initial models with PDB accession codes 7qkk and 6tjx (29). Atomic refinement was performed in ISOLDE using three rungs. Dihedral angles from the middle rung were set as a template for the rungs above and below. For each structure, Phenix refinement was performed of the first half-map. The resulting model was compared to this half-map and the second half-map to confirm the absence of overfitting (Supplementary Figure 1). Further details of model refinement, validation and statistics are given in Supplementary Table 1.

## AUTHOR INFORMATION

### Acknowledgements

We thank Jake Grimmett, Toby Darling and Ivan Clayson for help with high-performance computing.

### Funding

This work was funded by the Medical Research Council, as part of UK Research and Innovation (MC-UP-A025-1013 to S.H.W.S. and MC-U105184291 to M.G.). It was also supported by the Japan Agency for Science and Technology (Crest, JPMJCR18H3, to M.H.), the Japan Agency for Medical Research and Development (AMED, JP20dm0207072 and JP21wm0425019, to M.H.), The Japan Society for the Promotion of Science (JSJP KAKENHI, JP20K16482, to A.T.) and the Mochida Memorial Foundation for medical and Pharmaceutical Research (to A.T.).

### Author contributions

AT performed seeded aggregation, immunoblotting and immunoelectron microscopy; MH, ACR, DMAM, YS, SM and TT identified the patients and performed neuropathology; SL prepared cryo-EM samples and performed cryo-EM structure determination; XZ and AK acquired cryo-EM data; MG, SHWS and MH supervised the project and all authors contributed to the writing of the manuscript.

### Data availability

Cryo-EM maps have been deposited in the Electron Microscopy Data Bank (EMDB) with the accession numbers EMD-xxx, EMD-xxx. Corresponding refined atomic models have been deposited in the Protein Data Bank (PDB) under accession numbers xxx, xxx. Please address requests for materials to the corresponding authors. For the purpose of open access, the MRC Laboratory of Molecular Biology has applied a CC-BY public copyright licence to any Author Accepted Manuscript version arising.

### Competing interests

The authors declare that they have no competing interests.

## Supplementary Figures

**Supplementary Figure 1:**
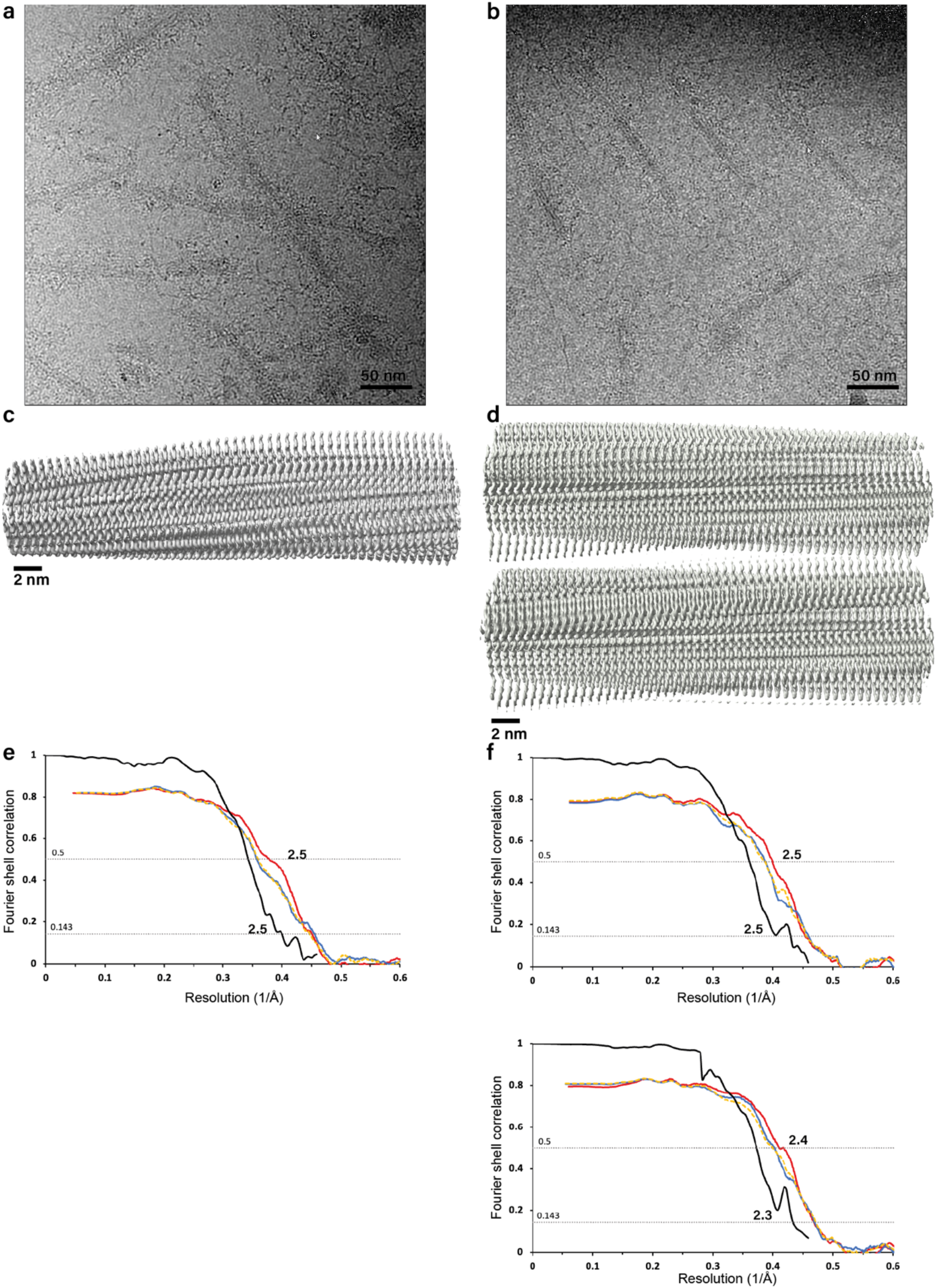
Cryo-EM structure determination. a-b. Representative micrographs of tau filaments from SH-SY5Y cells seeded with filaments from Alzheimer’s disease and corticobasal degeneration, respectively. c-d. Side views of reconstructed density of tau filaments from SH-SY5Y cells seeded with filaments from Alzheimer’s disease and corticobasal degeneration, respectively. e-f. Fourier shell correlation (FSC) curves for tau filaments from SH-SY5Y cells seeded with filaments from Alzheimer’s disease and corticobasal degeneration, respectively. Solvent-corrected FSC curves between independently refined half-maps are shown in black; FSC curves between the refined model and the reconstruction from all particles are shown in red; FSC curves between a model refined against half map 1 against half map 1 are shown in dashed yellow; FSC curves of the same model against half map 2 are shown in blue. In panels d and f, the seeded structure type 1 is shown above type 2.

**Supplementary Figure 2:**
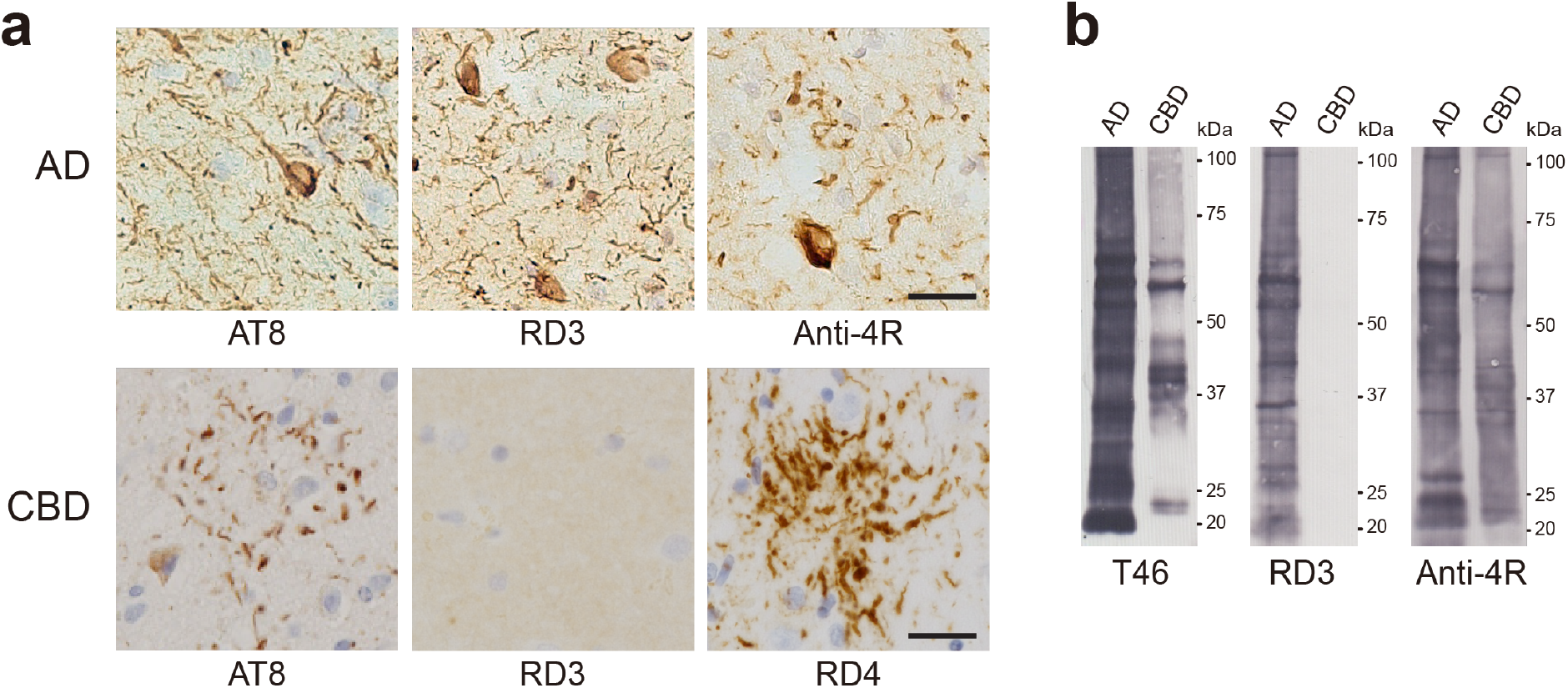
Imunohistochemistry and immunoblotting of AD and CBD cases. a. Immunostaining of sections from occipital cortex of the Alzheimer’s disease case and putamen of the corticobasal dewgeneration case using anti-tau antibodies AT8, RD3, Anti-4R and RD4. Scale bar, 25 mm. b. Immunoblotting of sarkosyl-insoluble fractions from occipital cortex of the Alzheimer’s disease case and putamen of the corticobasal degeneration case using anti-tau antibodies T46, RD3 and Anti-4R.

**Supplementary Figure 3:**
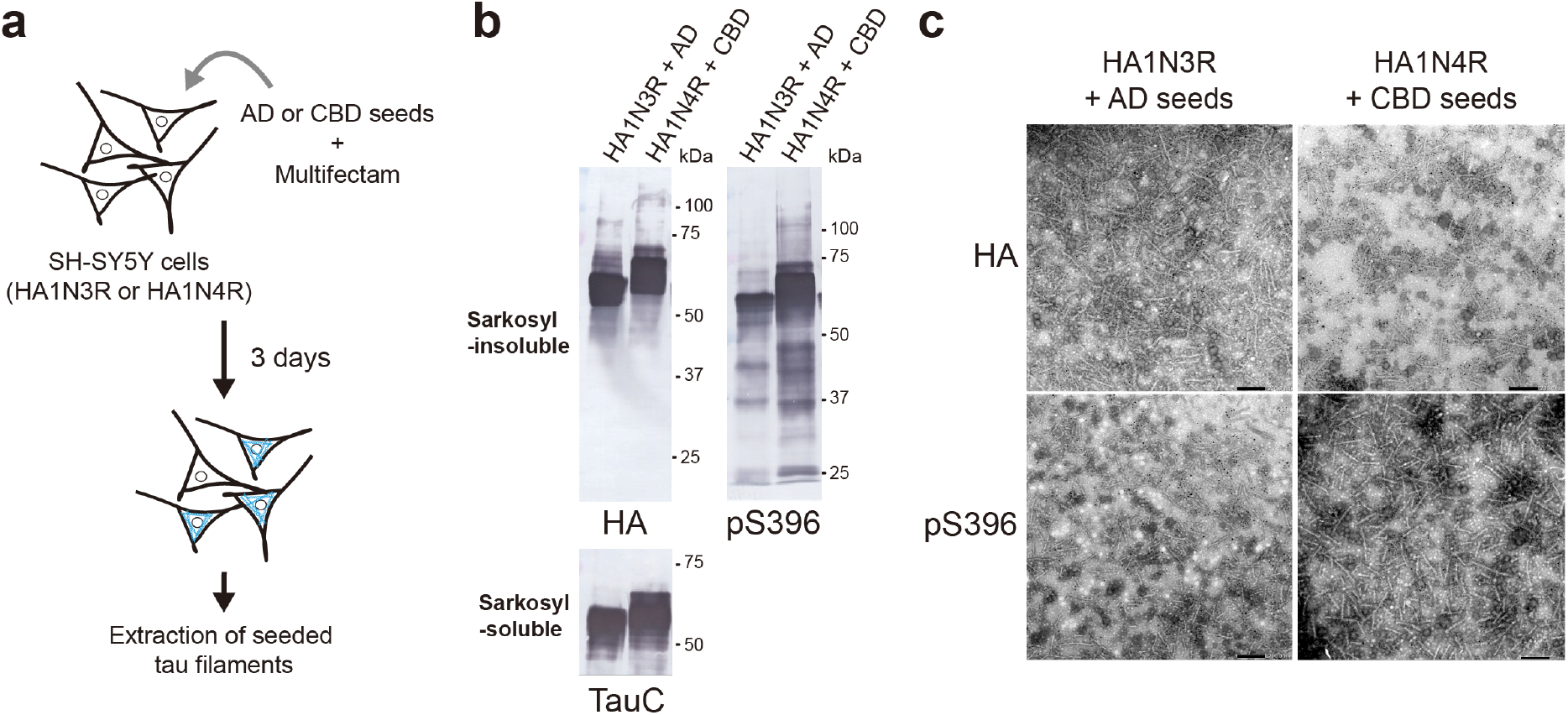
Immunoblotting and immunoelectron microscopy of tau assemblies from the sarkosyl-insoluble fractions of SH-SY5Y cells. a. Schematic of the seeded tau aggregation model. b. Immunoblotting of sarkosyl-insoluble tau from seeded SH-SY5Y cells using anti-HA and pS396 tau antibodies. Total tau was detected in the sarkosyl-soluble fraction using anti-tau antibody TauC. c. Immunoelectron microscopy of sarkosyl-insoluble tau frojm seeded SH-SY5Y cells using anti-HA and anti-pS396 tau antibodies. The secondary antibody was conjugated to 5 nm gold particles. Scale bar, 200 nm.

**Supplementary Table 1.**
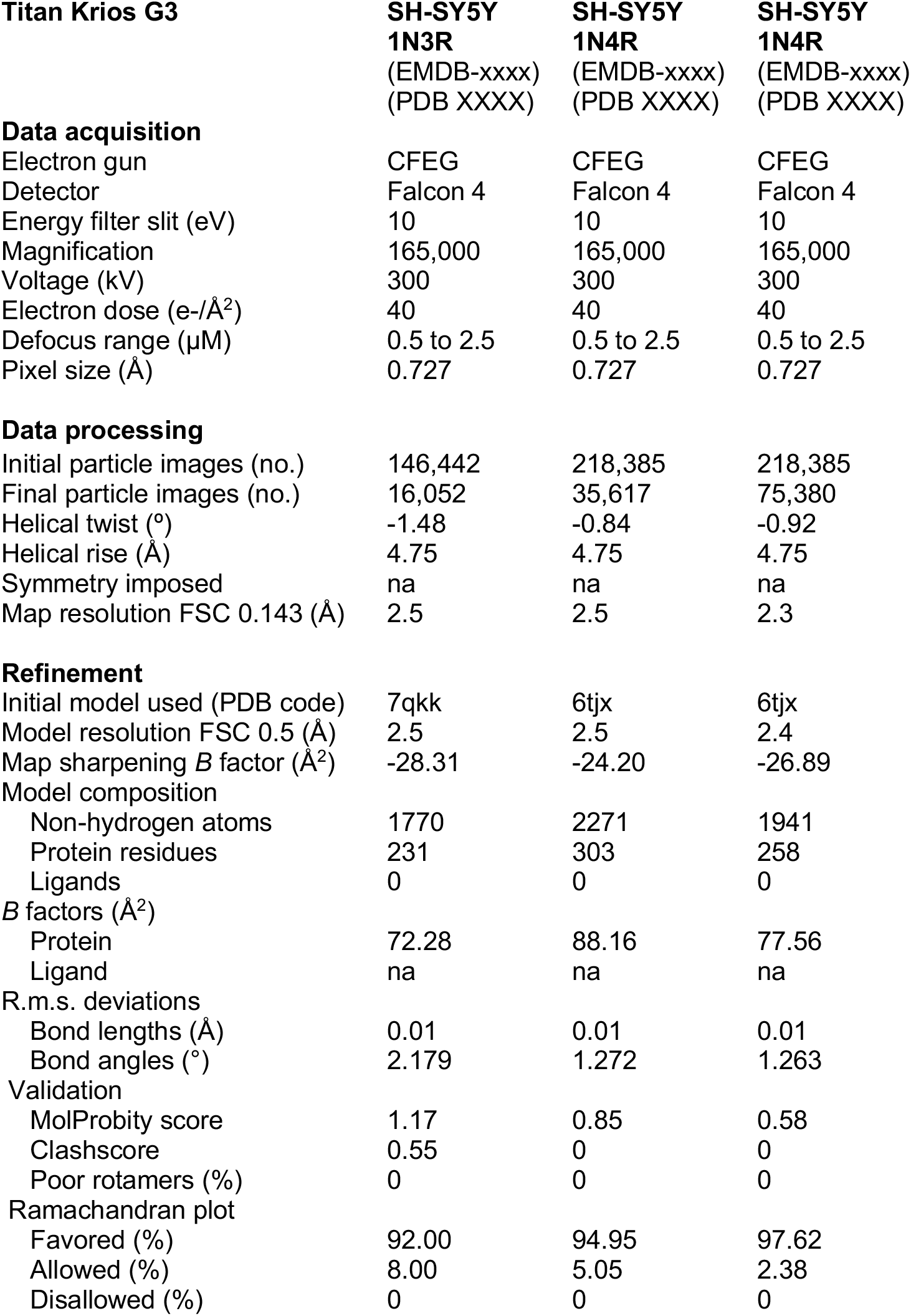

| **Titan Krios G3** | **SH-SY5Y**<br>**1N3R**<br>**(EMDB-xxxx)**<br>**(PDB XXXX)** | **SH-SY5Y**<br>**1N4R**<br>**(EMDB-xxxx)**<br>**(PDB XXXX)** | **SH-SY5Y**<br>**1N4R**<br>**(EMDB-xxxx)**<br>**(PDB XXXX)** |
| --- | --- | --- | --- |
| **Data acquisition** |  |  |  |
| Electron gun | CFEG | CFEG | CFEG |
| Detector | Falcon 4 | Falcon 4 | Falcon 4 |
| Energy filter slit (eV) | 10 | 10 | 10 |
| Magnification | 165,000 | 165,000 | 165,000 |
| Voltage (kV) | 300 | 300 | 300 |
| Electron dose (e-/Å<sup>2</sup>) | 40 | 40 | 40 |
| Defocus range (µM) | 0.5 to 2.5 | 0.5 to 2.5 | 0.5 to 2.5 |
| Pixel size (Å) | 0.727 | 0.727 | 0.727 |
| **Data processing** |  |  |  |
| Initial particle images (no.) | 146,442 | 218,385 | 218,385 |
| Final particle images (no.) | 16,052 | 35,617 | 75,380 |
| Helical twist (°) | -1.48 | -0.84 | -0.92 |
| Helical rise (Å) | 4.75 | 4.75 | 4.75 |
| Symmetry imposed | na | na | na |
| Map resolution FSC 0.143 (Å) | 2.5 | 2.5 | 2.3 |
| **Refinement** |  |  |  |
| Initial model used (PDB code) | 7qkk | 6tjx | 6tjx |
| Model resolution FSC 0.5 (Å) | 2.5 | 2.5 | 2.4 |
| Map sharpening *B* factor (Å<sup>2</sup>) | -28.31 | -24.20 | -26.89 |
| Model composition |  |  |  |
| Non-hydrogen atoms | 1770 | 2271 | 1941 |
| Protein residues | 231 | 303 | 258 |
| Ligands | 0 | 0 | 0 |
| *B* factors (Å<sup>2</sup>) |  |  |  |
| Protein | 72.28 | 88.16 | 77.56 |
| Ligand | na | na | na |
| R.m.s. deviations |  |  |  |
| Bond lengths (Å) | 0.01 | 0.01 | 0.01 |
| Bond angles (°) | 2.179 | 1.272 | 1.263 |
| Validation |  |  |  |
| MolProbity score | 1.17 | 0.85 | 0.58 |
| Clashscore | 0.55 | 0 | 0 |
| Poor rotamers (%) | 0 | 0 | 0 |
| Ramachandran plot |  |  |  |
| Favored (%) | 92.00 | 94.95 | 97.62 |
| Allowed (%) | 8.00 | 5.05 | 2.38 |
| Disallowed (%) | 0 | 0 | 0 |

## Notes

### Competing Interest Statement

The authors have declared no competing interest.

## References

1. Goedert M, Eisenberg DS and Crowther RA (2017) Propagation of tau aggregates and neurodegeneration. Annu Rev Neurosci 40, 189–210.

2. Goedert M, Spillantini MG, Jakes R, Rutherford D and Crowther RA (1989) Multiple isoforms of human microtubule-associated protein tau: Sequences and localization in neurofibrillary tangles of Alzheimer’s disease. Neuron 3, 519–526.

3. Fitzpatrick AWP, Falcon B, He S, Murzin AG, Murshudov G, Garringer HJ, Crowther RA, Ghetti B, Goedert M and Scheres SHW (2017) Cryo-EM structures of Tau filaments from Alzheimer’s disease brain. Nature 547, 185–190.

4. Falcon B, Zivanov J, Zhang W, Murzin AG, Garringer HJ, Vidal R, Crowther RA, Newell KL, Ghetti B, Goedert M et al. (2019) Novel tau fold in chronic traumatic encephalopathy encloses hydrophobic molecules. Nature 568, 420–423.

5. Falcon B, Zhang W, Murzin AG, Murshudov G, Garringer HJ, Vidal R, Crowther RA, Ghetti B, Scheres SHW and Goedert M (2018) Structures of filaments from Pick’s disease reveal a novel tau protein fold. Nature 561, 137–140.

6. Arakhamia T, Lee CE, Carlomagno Y, Duong DM, Kundinger SR, Wang K, Williams D, DeTure M, Dickson DW, Cook SN et al. (2020) Posttranslational modifications mediate the structural diversity of tauopathy strains. Cell 180, 633–644.

7. Zhang W, Tarutani A, Newell KL, Murzin AG, Matsubara T, Falcon B, Vidal R, Garringer HJ, Shi Y, Ikeuchi T et al. (2020) Novel tau filament fold in corticobasal degeneration. Nature 580, 283–287.

8. Shi Y, Zhang W, Yang Y, Murzin AG, Falcon B, Kotecha A, van Beers M, Tarutani A, Kametani F, Garringer HJ et al. (2021) Structure-based classification of tauopathies. Nature 598, 359–363.

9. Frost B, Jacks RL and Diamond MI (2009) Propagation of tau misfolding from the outside to the inside of a cell. J Biol Chem 284, 12845–12852.

10. Clavaguera F, Bolmont T, Crowther RA, Abramowski D, Frank S, Probst A, Fraser G, Stalder AK, Beibel M, Staufenbiel M et al. (2009) Transmission and spreading of tauopathy in transgenic mouse brain. Nature Cell Biol 11, 909–913.

11. Falcon B, Zhang W, Schweighauser M, Murzin AG, Vidal R, Garringer HJ, Ghetti B, Scheres SHW and Goedert M (2018) Tau dfilaments from multiple cases of sporadic and inherited Alzheimer’s disease adopt a common fold. Acta Neuropathol 136, 699–708.

12. Shi Y, Murzin AG, Falcon B, Epstein A, Machin J, Tempest P, Newell KL, Vidal R, Garringer HJ, Sahara N et al. (2021) Cryo-EM structures of tau filaments from Alzheimer’s disease with PET ligand APN-1607. Acta Neuropathol 141, 697–708.

13. Lövestam S, Schweighauser M, Matsubara T, Murayama S, Tomita T, Ando T, Hasegawa K, Yoshida M, Tarutani A, Hasegawa M et al. (2021) Seeded assembly *in vitro* does not replicate the structures of α-synuclein filaments from multiple system atrophy. FEBS Open Bio 11, 999–1013.

14. Tarutani A, Miyata H, Nonaka T, Hasegawa K, Yoshida M, Saito Y, Murayama S, Robinson AC, Mann DMA, Tomita T et al. (2021) Human tauopathy-derived tau strains determine the substrates recruited for templated amplification. Brain 144, 2333–2348.

15. Hallinan GI, Rejaul Hoq M, Ghosh M, Vago FS, Fernandez A, Garringer HJ, Vidal R, Jiang W and Ghetti B (2021) Structure of tau filaments in prion protein amyloidosis. Acta Neuropathol 142, 227–241.

16. Goedert M, Spillantini MG, Cairns NJ and Crowther RA (1922) Tau proteins of Alzheimer paired helical filaments: Abnormal phosphorylation of all six brain isoforms. Neuron 8, 159–168.

17. Lövestam S, Koh FA, van Knippenberg B, Kotecgha A, Murzin AG, Goedert M and Scheres SHW (2022) Assembly of recombinant tau into filaments identical to those of Alzheimer’s disease and chronic traumatic encephalopathy. eLife 11, e76494.

18. Braak H and Braak E (1991) Neuropathological stageing of Alzheimer-related changes. Acta Neuropathol 82, 239–259.

19. Kouri N, Whitwell JL, Josephs KA, Rademakers R and Dickson DW (2011) Corticobasal degeneration: a pathologically distinct 4R tauopathy. Nature Rev Neurol 7, 263–272.

20. Ling H, Kovacs GG, Vonsattel JPG, Davey K, Mok KY, Hardy J, Morris HR, Warner TT, Holton JL and Revesz T (2016) Astrogliopathy predominates the earlies stage of corticobasal degeneration pathology. Brain 139, 3237–3252.

21. Taniguchi-Watanabe S, Arai T, Kametani F, Nonaka T, Masami-Suzukake M, Tarutani A, Murayama S, Saito Y, Arima K, Yoshida M et al. (2016) Biochemical classification of tauopathies by immunoblot, protein sequence and mass spectrometric analyses of sarkosyl-insoluble and trypsin-resistant tau. Acta Neuropathol 131, 267–280.

22. Biedler JL and Schachner M (1978) Multiple neurotransmitter synthesis by human neuroblastoma cell lines and clones. Cancer Res 38, 3751–3757.

23. Zivanov J, Nakane T and Scheres SHW (2019) A Bayesian approach to beam-induced motion correction in cryo-EM single particle analysis. IUCrJ 6, 5–17.

24. Rohou A and Grigorieff N (2015) CTFFIND4: Fast and accurate defocus estimation from electron micrographs. J Struct Biol 192, 216–221.

25. He S and Scheres SHW (2017) Helical reconstruction in RELION. J Struct Biol 198, 163–176.

26. Scheres SHW (2020) Amyloid structure determination in RELION-3.1. Acta Crystallogr D 76, 94–101.

27. Zivanov J, Otón J, Ke Z, von Kügelgen A, Pyle E, Qu K, Morado D, Castaño-Díez D, Zanetti G, Bharat TAM et al. (2022) A Bayesian approach to single-particle electron cryo-tomography in RELION-4.0. eLife 11, e83724.

28. Scheres SHW and Chen S (2012) Prevention of overfitting in cryo-EM structure determination. Nature Meth 9, 853–854.

29. Casañal A, Lohkamp B and Emsley P (2020) Current developments in Coot for macromolecular model building of electron cryo-microscopy and crystallographic data. Protein Sci 29, 1069–1078.

